# Validating a screening agar for linezolid-resistant enterococci

**DOI:** 10.1101/798983

**Authors:** Guido Werner, Carola Fleige, Ingo Klare, Robert E. Weber, Jennifer K. Bender

## Abstract

Linezolid is an alternative treatment option for infections with multidrug-resistant Gram-positive bacteria including vancomycin-resistant enterococci (VRE). Some countries report an increasing number of isolates with resistance to linezolid. The recent publication of the Commission for Hospital Hygiene in Germany on enterococci/VRE recommends screening for linezolid-resistant enterococci (LRE). However, a suitable selective medium or a genetic test is not available. Our aim was to establish a selective screening agar for LRE detection and validate its application with a comprehensive collection of clinical LRE and linezolid-susceptible enterococci (LSE). We decided to combine the selective power of an enterococcal screening agar with a supplementation of linezolid. Several rounds of analyses with reference, control and test strains pointed towards Enterococcosel agar and a concentration of 2 mg/L linezolid. Finally, we validated our LRE agar with 400 samples sent to our National Reference Centre in 2019.

## Background

Linezolid is considered as one of the few remaining treatment options for infections with vancomycin-resistant enterococci (VRE) and other multidrug-resistant Gram-positive bacteria such as methicillin-resistant *Staphylococcus aureus* (MRSA) and/or methicillin-resistant *Staphylococcus epidermidis* (MRSE). The National Reference Centre (NRC) for Staphylococci and Enterococci recognized a growing number of linezolid-resistant enterococci (mainly *E. faecium*) and staphylococci (mainly *S. epidermidis*) from clinical samples in Germany in recent years (1, 2). The recent expiration of patent protection might have further promoted the more frequent and less critical use of linezolid in clinical practice. The association between the amount of linezolid usage and selection and detection of linezolid-resistant enterococci (LRE) and staphylococci has been addressed in several studies (3, 4). Also, a linezolid-dependent growth adaptation was described just recently (5). In accordance with rules of good antibiotic stewardship, a number of hospitals in Germany already reduced the use of linezolid and comparator substances or put their administration under specific internal clearance procedures, thereby limiting selective pressure and preserving the efficacy of this important last resort therapeutic for the most critical cases (6).

In 2018, the German Commission for Hospital Hygiene and Infection Prevention (“Kommission für Krankenhaushygiene und Infektionsprävention” - KRINKO) released a recommendation for the prevention of infections with “enterococci harboring special resistances” (7). This national directive focused not only on vancomycin resistance as the key resistance trait in clinical enterococci but also addressed the growing problem of vancomycin-resistant (and vancomycin-susceptible) enterococci with resistances to last resort antibiotics such as linezolid, tigecycline and daptomycin. As a recommendation, isolates with corresponding resistances or non-susceptibilities, especially to linezolid, should be handled similar to VRE. The guideline suggests screening for such isolates in case of supposed transmission events, e.g. when more than single cases are notified within three months and an epidemiological link between such isolates is expected. However, the technical implementation of this recommendation is less clear, since commercial agar media for the detection of linezolid or other last resort-resistances in enterococci are not yet available. Wildtype susceptible isolates and isolates categorized as “linezolid-resistant” differ by 1 – 2 dilution steps mostly, thus complicating a fine-tuning of supplementation of media with corresponding antibiotics. In the present study we tested different enterococcal agar media, supplemented with varying concentrations of linezolid, to determine the best media-antibiotic combination for reliable LRE screening.

## Materials and methods

All strains included in this study were received by the NRC as part of the routine work. Linezolid resistance was confirmed by broth microdilution according to EUCAST v9.0 and partly by a second, independent method (Etest® linezolid, bioMeriéux, Nürtingen, Germany). The isolates were genetically characterized for harboring 23S rDNA mutations associated with linezolid resistance and/or linezolid resistance genes such as *cfr*(B), *optrA* and *poxtA* (see later; **Supplementary Table S1**).

Pre-tests were performed with 3 commercially available agar media (i) Mueller-Hinton (Becton-Dickinson, Heidelberg, Germany), Enterococcosel Agar® (ECSA; Becton-Dickinson; Order No. 254019) and Bile-Esculin-Azide Agar (BEAA; Sigma-Aldrich, St. Louis, USA; Order No. 06105). Reference isolates *E. faecalis* ATCC 29212 (linezolid-susceptible; LIN MICs 1 – 4 mg/L), *E. faecium* ATCC 19434 (linezolid-susceptible; LIN MICs 1 – 2 mg/L), *S. aureus* ATCC 25923 (linezolid-susceptible) and *E. coli* ATCC25922 as well as five *E. faecium* and three *E. faecalis* isolates with linezolid MICs of 4 to >32 mg/L served as negative and positive control isolates, respectively (Table 1). The following procedure was applied to all tests except where specified differently: Microbial colonies were suspended in 4 ml Brain Heart Infusion broth and grown for 2h at 37 °C until an OD_650_ of 0,10 – 0,13 was reached. The suspension was diluted 1:10 in saline and 10 µl were plated onto the prepared selective agar media. Plates were incubated for 24 – 48h at 35 – 37°C. As a first step, all ten reference and clinical enterococcal strains (Table 1) were put onto (i) MH agar, (ii) ECSA and (iii) BEA agar supplemented with linezolid (Sigma-Aldrich) concentrations of 1, 2, 4, 8, 16, 32, 64, and 128 mg/L performed as dot blot experiments in order to narrow the linezolid test range. We repeated these experiments with the three agar brands supplemented with linezolid concentrations of 0, 1, 2, and 4 mg/L by streaking out the 10 µl of bacterial dilutions and by performing mixed culture growth experiments with (a) *E. coli* ATCC25922 / *S. aureus* ATCC25923 / *E. faecium* UW19369 (linezolid MIC = 32 mg/L) and (b) *E. coli* ATCC25922 / *S. aureus* ATCC25923 / *E. faecalis* UW17810 (linezolid MIC = 32 mg/L) in the same manner except for the *E. coli* and *S. aureus* isolates which were diluted 1:100 to reach similar colony counts as compared to the enterococcal strains.

**Table 1.** Linezolid MICs of two reference isolates and eight clinical strains on Mueller-Hinton, Enterococcosel and Bile Esculin Azid Agar measured after 24h and 48h incubation at 37 °C.

| Isolate | Species | Microdilution<br>LIN-MIC | MICs after 24h incubation on |  |  | MICs after 48h incubation on |  |  |
| --- | --- | --- | --- | --- | --- | --- | --- | --- |
|  |  |  | MHA | ECSA | BEAA | MHA | ECSA | BEAA |
| ATCC 29212 | *E. faecalis* | 1 – 4 | ≤1 | ≤1 | ≤1 | 2 | 2 | 2 |
| ATCC19434 | *E. faecium* | 1 – 2 | ≤1 | ≤1 | ≤1 | 2 | 2 | 2 |
| UW19506 | *E. faecium* | 8 | 4 | 2 – 4 | 2 – 4 | 16 | 8 | 4 – 8 |
| UW19369 | *E. faecium* | 32 | 8 | 2 – 4 | 2 – 4 | 32 | 8 | 8 – 16 |
| UW17810 | *E. faecalis* | 32 | 8 | 4 | 4 | 16 – 32 | 8 | 8 – 16 |
| UW17918 | *E. faecium* | 4 | ≤1 | ≤1 | ≤1 | 1 – 2 | 2 | 2 |
| UW19545 | *E. faecalis* | 4 | ≤1 | ≤1 | ≤1 | 1 – 2 | 2 | 2 |
| UW19507 | *E. faecium* | 16 | 8 | 2 – 4 | 2 | 8 – 16 | 8 | 4 – 8 |
| UW19236 | *E. faecium* | >32 | 16 – 32 | 4 – 8 | 4 | 32 | 16 – 32 | 16 |
| UW14781 | *E. faecalis* | 32 | 16 – 32 | 8 – 16 | 16 | 32 | 16 – 32 | 16 |

Legend: LIN, linezolid; MHA, Mueller-Hinton Agar; ECSA, Enterococcosel Agar; BEAA, Bile Esculin Acid Agar.

According to the results of the different pre-tests, extended analyses were performed with a single selective agar brand only and linezolid concentrations of a smaller range of 0, 2, and 3 mg/L. We included 48 test strains with linezolid MICs of ≤4 mg/L (susceptible; n= 6) and ≥8 mg/L (resistant; n= 42; **Supplementary Table S1**). The isolates originated from 23 diagnostic laboratories sent to the NRC in the first quarter of 2019. The strain collection was genetically diverse including linezolid-resistant strains harboring 23S rDNA mutations only and/or linezolid resistance genes such as *cfr*(B), *optrA* and *poxtA* (see later; **Supplementary Table S1**).

Finally a feasibility study was performed with the agar medium and the elaborated linezolid concentration as deduced from previous tests. Altogether 400 enterococcal isolates sent to the NRC between February and June 2019 as samples were put directly on linezolid screening agars with a fixed linezolid concentration (see Results). Plates were incubated at 37°C with readout after 24h and 48h.

Genomic DNA was prepared using the DNeasy Blood and Tissue Kit (Qiagen, Hilden, Germany) according to the manufacturer’s instructions. As an exception, cells were treated initially for 30 min at 37°C with lysozyme to achieve cell wall lysis. Genetic mutations in 23S rDNA alleles associated with linezolid resistance were determined by a procedure described earlier (8). The presence of mobile linezolid resistance genes *cfr*(B), *optrA* and *poxtA* was confirmed by a multiplex PCR as described recently (9).

Statistical calculations for sensitivity and specificity were carried out according to: https://www.medcalc.org/calc/diagnostic_test.php.

## Results

### Pre-tests to identify the optimal agar medium and linezolid test range

We have performed pre-tests with three media including a non-selective MH agar and two selective agar media, ECSA and BEAA. The agar media were supplemented with linezolid concentrations of 1 – 128 mg/L and growth of ten enterococcal reference and control strains was compared to growth on linezolid-free agar (see Methods; not shown in details). Linezolid MICs derived from agar dilution were 1- to 2-fold lower compared to the broth microdilution MICs (after 20h readout, Table 1). The agar dilution linezolid MICs increased generally by one step after 48h readout and were then in the similar range (+/- one dilution step) as compared to the broth microdilution linezolid MICs (Table 1). We did not see any agar-specific influence on the linezolid MIC.

We additionally performed mixed culture experiments with isolates of E.coli ATCC 25922, *S. aureus* ATCC 25923 and a linezolid-resistant strain of *E. faecalis* UW17810 and *E. faecium* UW19369, respectively (Table 1). Mixed cultures were applied to MHA, ECSA and BEAA supplemented with 0, 1, 2, and 4 mg/L linezolid and incubated for up to 48h. Black shades on ECSA and BEAA demonstrated growth of the enterococcal isolates, whereas growth on MHA might indicate the presence of both, *E. coli* and the corresponding LRE strain (not shown in details). Growth after 48h was visible on all tested ECSA and BEAA plates whereas after 24h black shaded growth was only visible on ECSA and BEAA plates with 1 mg/L linezolid (not shown in details).

Bacterial colonies grew larger on ECSA than on BEAA and thus this medium was selected for further experiments.

### Test series to identify the ideal linezolid concentration to screen for LRE

Results of all pre-tests (Table 1) revealed ECSA agar with a linezolid range between 2 to 4 mg/L to identify growth of LRE with a linezolid MIC of >4 mg/L. We tested a strain collection of 48 isolates (7 *E. faecalis*, 41 *E. faecium*) of which 42 were linezolid-resistant in broth microdilution (**Supplementary Table S1**). The linezolid MICs of these 48 isolates plus the two linezolid-susceptible reference isolates *E. faecalis* ATCC 29212 and *E. faecium* ATCC 19434 were measured on ECSA supplemented with 0, 2, and 3 mg/L linezolid after 24h and 48h readout. In general, growth was weak after 24 hours of incubation; hence most LREs could not be detected regardless of their linezolid MIC. Readout after 48h substantially increased the detection limit and visible growth of 35 isolates with linezolid MICs ≥8 mg/L was observed. The eight linezolid-susceptible wildtype and reference isolates did not grow on any plate supplemented with linezolid of 2 or 3 mg/L and incubated for up to 48h. Altogether six *E. faecium* isolates with a linezolid MIC of 8 mg/L in broth microdilution did not grow as well. A repeated phenotypic and molecular analysis of the latter six isolates revealed (i) microdilution MICs for linezolid of ≤2 to 4 mg/L for five isolates (1 isolate with 8 mg/L); a linezolid Etest MIC of 1.5 to 4 mg/L; (iii) no detectable 23S mutation and no presence of either *optrA* or *poxtA* (**Supplementary Table S1**). Linezolid resistance mutations and/or presence of resistance genes *optrA* or *poxtA* were also analyzed for all 35 isolates showing growth on selective ECSA. Seven isolates contained *optrA* (mostly *E. faecalis*), one *E. faecium* harbored *poxtA* and none of those eight isolates displayed any 23S rDNA mutation. All but one of the other isolates demonstrated 23S ribosomal mutations as the probable cause of linezolid resistance. The single exceptional isolate that only grew weekly on the selective agar did not contain any ribosomal rDNA or protein (*rplC, rplD*) mutation or resistance gene (*poxtA, optrA*), but revealed repeatedly resistant linezolid MIC in broth microdilution (8 – 16 mg/L) or Etest (12 mg/L). Based on these data we recommend a concentration of 2 mg/L linezolid supplemented with ECSA.

### Feasibility study to directly screen for LRE

The NRC does not receive original clinical samples but pre-characterized and pre-defined clinical isolates for further detailed analysis. We performed a feasibility study with 400 samples sent to the NRC within a 5-months’ time span in 2019. Altogether, 56 of the 400 samples received demonstrated linezolid MICs of >4 mg/L in subsequent broth microdilution assays. Readout after initial overnight incubation on LRE agar was almost impossible since most of these samples demonstrated only slight growth with grey shading but no single grown colonies. However, after 48h, a total of 54 out of 56 isolates grew on the LRE selective agar. The two isolates that did not grow on LRE agar after 48h were further analyzed. UW19813 revealed an Etest MIC of 4 mg/L but a microdilution MIC of 8 mg/L. Detailed analysis identified two colony morphologies when inspecting the original swab sample, but both supposedly different strains revealed an identical MIC profile in our panel of 18 different antibiotics. The other swab sample revealed growth after a repeated testing on LRE agar (isolate UW20075 with linezolid MICs 8 – 16 mg/L).

Altogether 26 additional samples demonstrated growth after 48h readout on LRE agar, but isolates later revealed linezolid MICs of ≤2 to 4 mg/L (= susceptible). This means that 26 of 342 swab samples (7,6%) with *Enterococcus* spp. isolates with susceptible linezolid MICs in broth microdilution generated a false positive result (two swab samples did not contain *Enterococcus* spp. isolates at all and were completely excluded from further analyses and calculations). A repeated testing starting with these 26 original samples revealed the following: (i) 17 samples did not show colony growth on LRE agar and Etests of corresponding isolates revealed MICs of ≤4 mg/L (= susceptible); (ii) nine samples demonstrated colony growth again but (newly) isolated strains revealed linezolid MICs of >4 mg/L (= resistant). Six of the latter samples revealed more than one strain after repeated and detailed phenotypic analysis (“mixed cultures”) which might have complicated subsequent phenotypic and genetic analysis. Diagnostic test performance not considering repeated test results revealed a sensitivity of 96.4% (CI: 87.7 to 99.6%), a specificity of 92.4% (CI: 89.1 to 95%), a PPV of 67.5% (CI: 58.9 to 75.1%) and a NPV of 99.4% (CI: 97.6 to 99.8%). When original data were corrected for results of repeated and genetic confirmatory tests, all values increased leading to a sensitivity score of 98.4% (91.6 to 100%), specificity of 100% (CI: 98.9 to 100%), PPV of 100% and NPV of 99.7% (CI: 98.6 to 100%).

## Discussion

In 2018 the German Commission for Hospital Hygiene KRINKO released a recommendation of how to deal with hospitalized patients colonized and infected with enterococci with special resistances including VRE. This national guideline recommended screening for enterococci with resistances to last resort antibiotics such as linezolid if clusters of more than one isolates infecting or colonizing patients within a 3-months-time span are noticed (7). However, the guideline did not recommend a specific diagnostic test to implement this screening procedure for LRE in daily laboratory routine, and to the best of our knowledge, such test or test medium was not available at the beginning of our study. Meanwhile, a recent analysis was published suggesting a Super Linezolid agar which is based on MH supplemented with linezolid of 1.5 mg/L (10). The authors additionally supplemented the agar with aztreonam (2 mg/L), colistin (15 mg/L) and amphotericin B (5 mg/L) to suppress microorganisms of the normal intestinal flora which would otherwise grow on the non-selective MH medium. Although the agar was designed to screen for linezolid-resistant Gram-positive cocci, the collection contained primarily linezolid-resistant *S. epidermidis* (n= 13), but only a very limited number of linezolid-resistant isolates of other genera and species such as *S. aureus* (n= 2), *S. capitis* (n= 1) or *E. faecium* (n= 1). The collection did not contain enterococcal isolates with low linezolid MICs fairly above the breakpoint of >4 mg/L as it is typical for gene-based linezolid resistance encoded by *optrA* or *poxtA* in *Enterococcus* spp. (see **Suplementary Table S1**). Thus, the results of the Super Linezolid medium could hardly be compared with our study results which explicitly focused on a selection and identification of LRE.

Within the present study we combined the selective power of enterococcal screening agars with a supplementation of an ideal concentration of linezolid to screen for LRE in original swab samples. The two different enterococcal agar media analyzed within this study only showed minor differences in growth. We are well aware of further brands and producers which could not be included and tested all in this study (e.g., Bile Esculin Agar, MAST Diagnostika, Reinfeld, Germany; Bile Esculin Agar, Oxoid/Thermo-Fisher, Wesel, Germany; Bile-Esculin-Azid Agar, Roth GmbH, Karlsruhe, Germany). Composition of these media is comparable to the two brands tested in this study. When we argue for ECSA due to certain specificities this should not implement that other, alternative producers might not also perform comparably well. In fact, the fine-tuning of the ideal linezolid concentration seems much more important than the brand or producer. A MIC of 8 mg/L or higher implements sufficient growth of all resistant bacteria at a supplemented linezolid concentration of 4 mg/L; however, a substantial number of the herein described test series revealed 2 mg/L linezolid as the best compromise between sufficient sensitivity and specificity and already 3 mg/L linezolid led to a fairly reduced growth of LRE (**Supplementary Table S1**). In all assays 48h incubation was essential which, admittedly, is far less acceptable in daily laboratory and hospital routine. Since we don’t have a better test yet and we lack a reliable genetic test, the herein described assay with a 48h incubation time is the best we can currently recommend meeting diagnostic and infection prevention and control requirements.

Our study was limited by the fact, that our suggested medium was not tested with clinical samples such as rectal swabs or stool samples. Although the components of enterococcal screening agars such as high salt, sodium azide and bile acid concentrations suppress growth of many other intestinal microorganisms, we could only speculate about general performance of our suggested medium with the aforementioned original clinical samples. Demonstration of degradation of the supplemented esculin by enterococcal growth on ECSA and BEAA leading to black agar color and black colonies is an important additional diagnostic feature. We consider that low colony counts of LRE in original stool samples or rectal swabs might reduce performance as well as other constituents being able to degrade esculin and as such color the agar grey or black simulating enterococcal growth. The authors of the aforementioned study describing the Super Linezolid agar (containing 1.5 mg/L of linezolid) performed such analysis with spiked stool samples and reached a quite low detection limit, also for their LRE strain (10). Also, results of our mixed culture experiments were promising. We could easily identify LRE on our ECSA medium supplemented with linezolid whereas other components of the bacterial mix did not grow.

In summary, we explicitly tested and validated a screening agar for linezolid-resistant isolates of *E. faecium* and *E. faecalis*. We recommend using an enterococcal selective agar such as Enterococcoselagar or a similar brand adding a supplementation of 2 mg/L linezolid. Growth of single colonies in combination with a black colony color after 48h incubation is indicative of a LRE.

## Acknowledgements

We thank Christine Günther and Karsten Großhennig for excellent technical assistance. We are thankful to all collaborators of the National Reference Centre for Staphylococci and Enterococci sending us corresponding strains for further analyses. A special thanks is dedicated to Prof. Heike von Baum for critically reading and reviewing the manuscript.

## Ethical approval

None applicable here.

## Availability of data and materials

Detailed microbiological data are available in the Supplementary Table S1. Further information and all original data and strains are available on request to G.W..

## Author statements

G.W., J.K.B., R.E.W. and I.K. designed the study. C.F. performed most of the experiments. C.F. and G.W. evaluated the study results. G.W. wrote the manuscript; all other authors reviewed the manuscript.

